# Open-source design and methods for a wide use of ferrofluid droplets micro-rheology

**DOI:** 10.64898/2026.11.24.768889

**Authors:** Nicolas Harmand, Camille Curantz, Thomas Panier, Olga Vasiljevic, Marie Anne Breau, Lea-Laetitia Pontani

## Abstract

Mapping the mechanical properties of tissues and the stresses that are present in developing organisms is central to the understanding of many morphogenetic processes. To this end, droplet-based micro-rheology has been developed as an interesting tool because of the method’s biocompatibility and non-destructive nature. In particular, ferrofluid droplets offer the possibility to probe directionally, locally and over time the evolution of mechanical properties in a developing tissue. We propose a platform to allow a wide use of this technique and to extend its capabilities. First, we present an open-source design for a magnetic field generator that can be readily 3D printed and reversibly mounted on classical fluorescence microscopes. Second, the ferrofluid is commercially available, yet biocompatible on experimental timescales. Finally, we explore variations of the field application protocols to show how this technique can probe a range of tissue properties and to evidence plastic behaviours. We apply these methods in four locations of zebrafish embryos that illustrate the great variety of properties and underlying biological processes that can be studied in developing tissues.

## INTRODUCTION

Morphogenesis is the process by which embryonic cells and tissues acquire their shape, organisation and spatial relationships through the coordinated integration of chemical and mechanical cues. During this process, the material properties of cells and tissues (their elasticity or stiffness, their viscosity and their plasticity) govern their ability to flow or deform in response to internal and external mechanical forces (Cao et al., 2026; Villedieu et al., 2020). Forces and the stiffness of the environment also act as direct mechanical cues, influencing cell behaviour and fate through mechanotransduction pathways that modify intracellular signalling and gene expression (Discher et al., 2005; Lo et al., 2000; Engler et al., 2006; Kechagia et al., 2019; Dupont and Wickström, 2022). In addition, tissue rigidity changes and force-induced morphogenetic movements can modify tissue architecture, thereby modulating the distribution and transduction of morphogens or other types of chemical cues (Fulton et al., 2022; Autorino et al., 2026). At the tissue scale, stiffness and mechanical forces contribute to morphogenesis but also homeostasis, wound healing and disease progression (Heisenberg and Bellaïche, 2013; Vining and Mooney, 2017; Mammoto and Ingber, 2010; Humphrey et al., 2014; Wozniak and Chen, 2009). Understanding these processes therefore requires determining the forces, or stresses, generated within tissues but also the material properties of the tissues themselves (Petridou and Heisenberg, 2019; Xi et al., 2019; Kayal et al., 2026). Indeed, biological tissues are mechanically complex, exhibiting combinations of elastic, viscous and plastic behaviours whose relative contributions can evolve over time (Cao et al., 2026; Tlili et al., 2015).

The mechanical *stress* is the force per unit area transmitted within the tissue. The *material properties* are defined by the relation between the stress a tissue carries and the resulting deformation, or strain. Rheological measurements are thus designed to establish that relation.

Elastic solids recover their initial shape when the stress is removed, and the relationship *σ* = *Eε* between a uniaxial stress *σ* and the resulting strain *ε* defines the Young’s modulus *E*, or modulus. In a viscous fluid the stress is set instead by the strain *rate*, 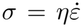, which defines the viscosity *η*, and the deformation accumulates for as long as the stress lasts. A viscoelastic material exhibits both properties, separated by a timescale *τ* ∼ *η/E*: a viscoelastic liquid deforms like a solid below *τ* and flows beyond, while a viscoelastic solid deforms progressively over *τ* and then holds a finite deformation. A plastic material retains an irreversible shape change once the stress has exceeded a yield stress. Biological tissues can display all these behaviours, which is why no single modulus describes them straightforwardly.

Several methods have been developed to quantify the mechanical properties of biological materials, each with a defined scope and scale: atomic force microscopy measures stiffness at high resolution but is restricted to an accessible surface (Krieg et al., 2019, 2008; Barriga et al., 2018; Pillai et al., 2026); micropipette aspiration returns a viscoelastic modulus or a surface tension of a whole cellular aggregate (Guevorkian et al., 2010; Hochmuth, 2000), or of single cells within an aggregate (Guevorkian and Maître, 2017); laser ablation and optical tweezers probe tensional forces at individual cell junctions (Rauzi et al., 2008; Bambardekar et al., 2015) or at the tissue scale (Collinet et al., 2015; Bonnet et al., 2012); and Brillouin microscopy provides a contact-free mechanical signal but does not lead to an absolute measurement of the elastic modulus (Scarcelli and Yun, 2008; Prevedel et al., 2019; Sánchez-Iranzo et al., 2020; Bevilacqua et al., 2019). To map out stresses within tissues, a second class of methods proposes to embed a calibrated deformable probe inside the tissue in order to infer stresses from its shape. These deformable probes can be either oil microdroplets (Campás et al., 2014; Vasiljevic et al., 2026), elastic or hydrogel microbeads (Dolega et al., 2017; Mohagheghian et al., 2018; Träber et al., 2019; Vorselen et al., 2020; Souchaud et al., 2022). Given that the material properties of the probes are known, their deformation under local stresses in the tissue allows them to be measured. For instance, liquid incompressible oil droplets are defined by their interfacial tension, which quantifies how much force needs to be applied to deform the droplets away from spherical. However, because these systems only deform passively, they can report mechanical stresses but do not allow micro-rheology experiments to be performed in order to measure the material properties of the tissues.

Using ferrofluid oil droplets overcomes this limitation. A biocompatible ferrofluid drop injected into the tissue acts as a passive stress sensor and, under a magnetic field, as an actuator (Campás et al., 2014; Serwane et al., 2017), allowing for local rheology measurements. Its diameter can be set from cellular to supracellular sizes, which defines the scale that is probed. With no applied field, its resting shape, knowing its interfacial tension, reports on the anisotropic stresses applied by the tissue just like for simple oil droplets. Under an applied homogeneous magnetic field, the magnetic moments of the nanoparticles dispersed in the oil align with the field, and the resulting magnetisation of the droplet elongates it into an ellipsoid along the field. This magnetic stress applied on the droplet is then balanced by its interfacial tension and by the resistance of the surrounding tissue: the equilibrium shape thus yields a modulus, while the dynamics of deformation and relaxation provide the characteristic viscoelastic timescales of the tissue (Serwane et al., 2017; Rowghanian et al., 2016; Bacri and Salin, 1982). This approach has been applied in pioneering work by the Campás lab, which showed that in zebrafish embryos, anteroposterior axis elongation is associated with a fluid-to-solid jamming transition in the tailbud region. The most posterior part, the progenitor zone, was shown to have a more fluid behaviour than the more anterior presomitic mesoderm (PSM) (Mongera et al., 2018), reflecting a gradient in the mechanics of the cellular microenvironment along the body axis in this area (Mongera et al., 2023; Stooke-Vaughan et al., 2025).

The active ferrofluid droplet has nonetheless seen limited deployment since its introduction, for reasons that may be practical rather than conceptual. Applying and controlling the magnetic field has so far required the assembly of a fixed set-up on a dedicated microscope: a permanent-magnet ring is translated towards and away from the sample to switch the field ON and OFF (Serwane et al., 2017), which occupies the instrument and sets the field direction once and for all. In addition to this, preparing the droplets means formulating and calibrating a ferrofluid, sometimes with dedicated chemical synthesis (Serwane et al., 2017). Finally, converting the recorded shapes into mechanical quantities takes a dedicated analysis and modelling framework.

Here we address these three barriers and propose a full system that is easily deployable across any lab. For the magnetic field, we use a removable module of two concentric, motorised Halbach rings of permanent magnets (Halbach, 1980; Hilton and McMurry, 2012; Tretiak et al., 2019): it mounts reversibly on a standard microscope, needs no power supply while imaging, hence avoiding heating, leaves the optical path clear, and is built from low-cost, 3D-printed parts for which we provide a construction blueprint. For the droplets, we use a commercial ferrofluid with a single added surfactant, so that no chemical synthesis is required. For the analysis, we provide an open-source pipeline that derives mechanical quantities from the recorded images; the droplet contour is measured on bright-field images alone, so no fluorescent labelling of the ferrofluid is needed, which frees every channel of the microscope for specific labelling. Because the two rings rotate independently, the field is *reconfigurable*: rotating them together sets its direction in the rings plane, and counter-rotating them sets its amplitude between zero and twice that of a single ring. Beyond an equilibrium stiffness, this gives access to the anisotropic residual stress of the tissue with its direction, to the relaxation times of the tissue, and to plasticity tests along distinct axes in the tissues obtained by rotating the field by 90^*°*^ relative to a pre-existing droplet deformation.

We calibrate the system in polyacrylamide gels of a composition whose Young’s modulus has been measured by bulk oscillatory rheometry (Cresens et al., 2026), then apply it in the developing head of the zebrafish embryo, where the brain, eye and olfactory placode tissues undergo extensive morphogenetic movements that have been shown to depend, at least in part, on mechanical cues and on mechanical interactions between neighbouring tissues (Moreno-Mármol et al., 2021; Sidhaye and Norden, 2017; Monnot et al., 2022; Breau et al., 2017; Tignard et al., 2024; Norden, 2023; Ramos et al., 2025). We finally apply the method to the zebrafish tailbud region, targeting the PSM, where published ferrofluid measurements exist and a direct comparison is possible.

## RESULTS

### A reconfigurable permanent-magnet field generator

Our method requires the application of a magnetic field of controlled amplitude and orientation on the biological samples. We took advantage of Halbach rings (Halbach, 1980), which consist of a circular arrangement of permanent magnets whose orientations are set to produce a spatially uniform field in the plane of their central bore. The amplitude of the field *B* is set by the ring geometry and the magnet strength, and its direction *θ*_*B*_ rotates with the ring. Nesting two such rings coaxially gives a net central field equal to the vector sum of the two (see Fig. 1B and Fig. S1). If the rings have equal strength and are set at angles *ϕ*_1_ and *ϕ*_2_, the field amplitude and direction yield

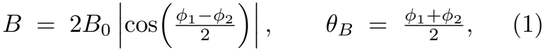

**FIG. 1.**
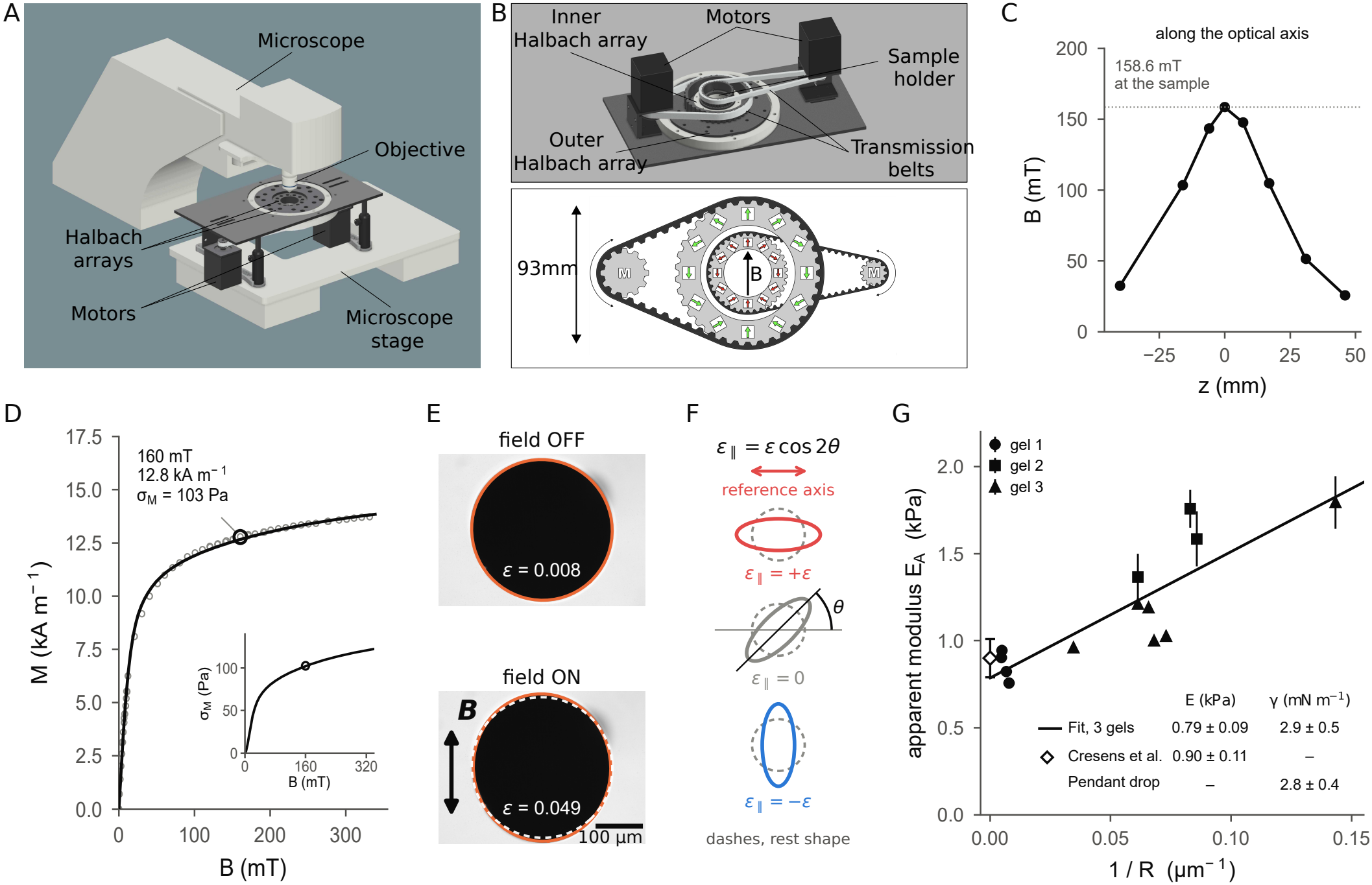
The reconfigurable field module and its calibration. (A) 3D view of the module fitted onto a microscope stand. (B) Top: upside down 3D view of the motorised module. Bottom: sketch of the module. Black arrow, magnetic field *B*, in the plane of the rings. Each ring is a *k* = 2 Halbach array containing twelve permanent magnets (red and green arrows indicating magnetisation direction). The bore is open, so the optical path is unobstructed. Each ring is driven by a stepper motor (labelled M) through a timing belt. Rotating the two rings in the same direction turns the field orientation, while counter-rotating them sets its amplitude between zero and twice that of a single ring. (C) Field characterisation: axial profile *B*(*z*) through the bore centre, measured with a Hall probe. The working point is 158.6 mT in the sample plane. (D) Magnetisation *M* (*B*) of the formulation used throughout (Tween 20 at 20 mg mL^*−*1^) measured by vibrating-sample magnetometry (points), with a Langevin fit (line). At the working field of 160 mT (open circle), *M* = 12.8 kA m^*−*1^ and *σ*_*M*_ ≃ 103 Pa. Inset, *σ*_*M*_ as a function of the field *B*. (E) Ferrofluid droplet in a polyacrylamide gel, *R* = 148 µm, with the field OFF and ON and the sub-pixel ellipse fit overlaid (orange). Black arrow, field direction; dashed contour, the field OFF shape reported on the field ON image. (F) Schematic of signed deformation measurements. The deformation is measured in the frame of the image, and its signed projection on a fixed reference axis is *ε*_∥_ = *ε* cos 2*θ*, with *θ* the angle between the droplet’s long axis and that reference. *ε*_∥_ is positive when the droplet is elongated along the reference axis, negative when along the orthogonal one, and zero at 45^*°*^. The reference is fixed and is not the current field axis, so a droplet aligned with a field applied along the orthogonal axis gives *ε*_∥_ *<* 0. Dashed lines: circular rest shape. (G) Apparent modulus *E*_*A*_ against 1*/R* for 13 droplets in three independent polyacrylamide gels of the same composition (6 % acrylamide, 0.06 % bis-acrylamide). Dark line: affine fit 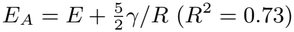 . Inset table: *E* and interfacial tension *γ* read from the linear fit, bulk modulus measured in Cresens et al. (2026) (open diamond), tension measured by pendant-drop experiments (Fig. S2). Error bars, measurement uncertainty of each droplet (see Materials and Methods).

so that the *half-difference* of the ring angles sets the amplitude, from 0 to 2*B*_0_, and the *half-sum* sets the direction. Two motors therefore give full and independent control of field amplitude and direction in the imaging plane. Because the field is produced by permanent magnets, the module needs no power supply, therefore no cooling and no current regulation while imaging, making it compact enough to be assembled into a portable device.

We operate at a working amplitude of 160 mT, measured as 158.6 mT on axis at the sample plane (Fig. 1C). The field is uniform to 2 *×* 10^*−*5^ % over a droplet, ensuring the droplet is only deformed and not moved from its position (see Materials and Methods).

### Droplet formulation

The droplets are made from a commercial ferrofluid oil (see Materials and Methods) supplemented with the biocompatible surfactant Tween 20 at 20 mg mL^*−*1^, which lowers the oil/water interfacial tension into the range where the deformations are optically measurable. Two quantities control their mechanics: the magnetisation *M* (*B*), which sets the stress we apply, and the interfacial tension *γ*, which sets the stress the droplet resists with.

The interfacial tension of the oil against phosphate-buffered saline (PBS) is measured by axisymmetric drop shape analysis on pendant drops (Fig. S2A) as a function of the Tween 20 concentration. This tension reaches *γ* = 2.79 *±* 0.37 mN m^*−*1^ at 20 mg mL^*−*1^ which is our working concentration throughout this article (see Fig. S2B).

We measured *M* (*B*) by vibrating-sample magnetometry (Dodrill and Lindemuth, 2021) on this formulation (Fig. 1D). This allows us to estimate the applied stress *σ*_*M*_ as a function of *B* through the relationship *σ*_*M*_ = *µ*_0_*M* ^2^*/*2, with *µ*_0_ the permeability of free space (see inset in Fig. 1D). At our typical working field of 160 mT, the magnetisation is 12.8 kA m^*−*1^, giving an applied magnetic stress *σ*_*M*_ = 103 Pa.

### From droplet shape to stress and stiffness

In the absence of an applied magnetic field, the resting shape of the droplet directly reports the residual anisotropic stress in the tissue, or pre-stress, *σ*_*T*_ ≃ (6*γ/R*) *ε*_off_, with *ε*_off_ = (*b − R*)*/R* the droplet strain in the absence of field, *b* being the semi-major axis of a fitted ellipse around the droplet, *R* the undeformed droplet radius and *γ* the droplet interfacial tension. In this case, the droplet can be used to map out stresses inside living tissues (Campás et al., 2014; Serwane et al., 2017; Vasiljevic et al., 2026).

In micro-rheology experiments, i.e. when a field is applied through the set-up, the relationship between droplet shapes and tissue mechanical properties can be derived by computing the balance between the magnetic stress, that tends to deform the droplet, and the mechanical properties of the droplet and surrounding tissue, that resist deformation, as done by Serwane et al. (2017) following Style et al. (2015). Four equations are needed here.

Given the formula of the droplet strain *ε* = (*b − R*)*/R*, if the droplet is considered as a prolate spheroid of semiaxes *b, a, a*, with *a* the semi-minor axis, volume conservation gives *b a*^2^ = *R*^3^ with *R* the rest radius, i.e. (*b/a*)^2*/*3^ = *b/R*, and the strain can be written as

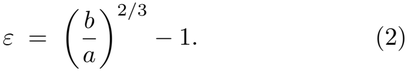

Since only the ratio *b/a* enters Eq. (2), *ε* does not need any spatial calibration. From each bright-field image the pipeline measures the orientation *θ* of the droplet’s long axis in the field of view. We fix a reference frame in the image and use the signed projection of the strain on the *x* axis, *ε*_∥_ = *ε* cos 2*θ* with *θ* measured from that axis: it is positive when the droplet is elongated along the reference axis, negative when along the orthogonal one, and zero at 45^*°*^ (Fig. 1F). The reference axis is fixed rather than tied to the current field direction, so a droplet deformation along a field applied in the orthogonal axis gives *ε*_∥_ *<* 0.

A droplet in a purely viscous medium is restored to a sphere by its own interfacial tension, and the balance against the applied magnetic stress reads

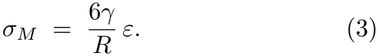

In an elastic medium of Young’s modulus *E* the droplet must in addition strain the surrounding solid, which adds a second elastic component. Both contributions can be modelled as two springs in parallel in classical rheology models (Fig. S3A), leading to

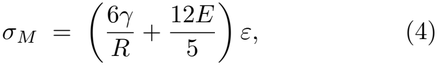

which is the incompressible limit of the liquid-inclusion problem (Eshelby, 1957; Style et al., 2015). This equation can be written as 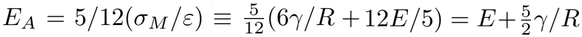, which means that the Young’s modulus *E* and the interfacial tension *γ* can be extracted by plotting *E*_*A*_, i.e. 5*/*12(*σ*_*M*_ */ε*), against 1*/R*.

Finally, in case the droplet is deformed by a pre-stress *σ*_*T*_ in the absence of a field, both *σ*_*M*_ and *σ*_*T*_ act on the droplet in the presence of a field. The difference between the deformations with and without field then leads to:

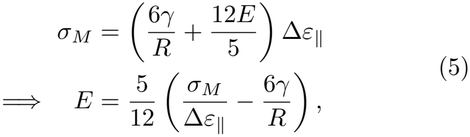

with Δ*ε*_∥_ the field-induced modulation of the *signed projected* strain.

### Calibration in polyacrylamide gels

To calibrate the system, we prepared three independent polyacrylamide gels of a single composition, 6 % acrylamide and 0.06 % bis-acrylamide, whose Young’s modulus measured by bulk rheometry is *E* = 0.90 *±* 0.11 kPa (Cresens et al., 2026), comparable to the stiffness reported in the zebrafish tailbud at short timescales, 400 Pa to 800 Pa (Mongera et al., 2023) (see Fig. 1E,G and Materials and Methods). Thirteen droplets of radii ranging from 7 µm to 212 µm were injected and probed across the three gels (see Materials and Methods). Each droplet was imaged with the field OFF, then with the field ON.

According to Eq. (4), fitting *E*_*A*_ against 1*/R* leads to an experimental estimate of the gel’s Young’s modulus at the intercept with the y-axis, while the interfacial tension *γ* is given by the slope.

The fit carried out on the three gels together (Fig. 1G) gives *γ* = 2.9 *±* 0.5 mN m^*−*1^ between the oil and the gel, consistent with the value measured by pendant drop against PBS, 2.79 *±* 0.37 mN m^*−*1^, and an intercept *E* = 0.79 *±* 0.09 kPa, within the uncertainty of bulk rheometry measurements (Cresens et al., 2026).

### Measurement of anisotropic stresses across tissues

We microinjected droplets in the tailbud region of zebrafish embryos, targeting the PSM, and imaged them at the 6–10 somite stage (Fig. 2A), 12–14 hours post fertilisation (hpf) (Kimmel et al., 1995); this region has already been studied with a similar method (Mongera et al., 2018). We also injected droplets at three locations of the developing head (Fig. 2B), at later stages (18–22 somites, 18–20 hpf): inside the eye, inside the olfactory placode, and at the interface between these two tissues, as their mechanical coupling through the extracellular matrix (ECM) has been reported to play a role in the morphogenesis of the olfactory placodes (Monnot et al., 2022; Breau et al., 2017). For droplet shape analysis we used bright-field imaging as their contrast is sufficient to visualise them and find their contour (Fig. 2C). Confocal imaging of embryos with labelled membranes localised the droplets and showed no detectable toxicity for the cells in direct contact with them, suggesting that our ferrofluid formulation is biocompatible on these experimental timescales (Fig. 2D and Materials and Methods).

**FIG. 2.**
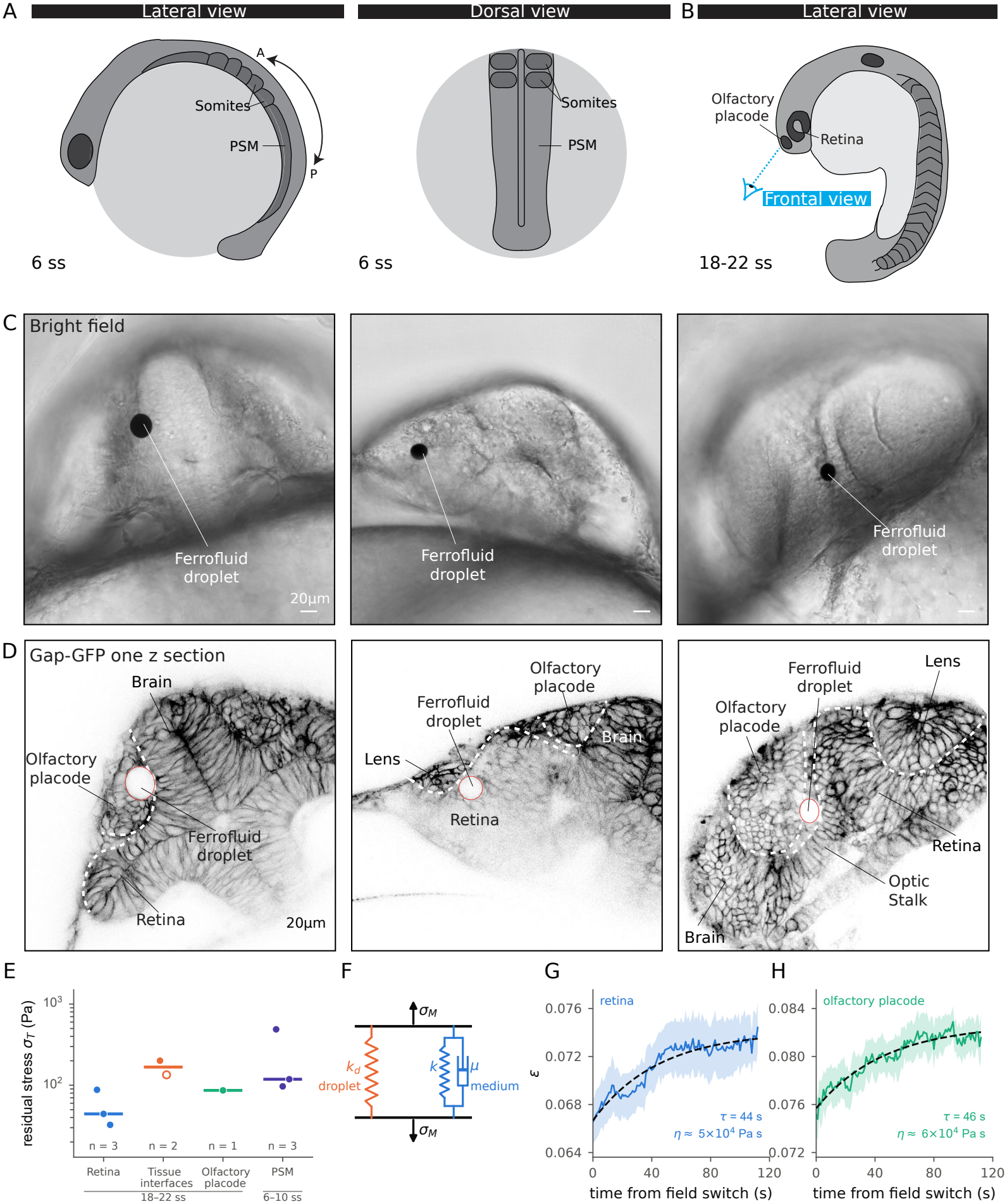
Droplets in the zebrafish head and tailbud: localisation, residual anisotropic stress and field response. (A) Cartoons showing lateral (left) and dorsal (right) views of a zebrafish embryo at the 6-somite stage (ss). (B) Cartoon of a lateral view of a zebrafish embryo at 18–22 ss; the blue line marks the frontal view of panels C and D. (C) Bright-field images showing field OFF droplets at the three different locations in the heads of zebrafish embryos at 18–22 ss, frontal (left) and fronto-lateral views (middle and right). (D) Corresponding confocal images of wild-type zebrafish embryos injected with gap-GFP mRNA, which labels all cell membranes. Displayed in inverted LUT, membranes appear in black and the ferrofluid droplets in white (outlined by a red circle) in the olfactory placode, in the eye and at the eye/olfactory placode interface. Dashed white line shows the outline of different head tissues. (E) Residual anisotropic stress *σ*_*T*_ = 6*γε*_off_ */R* read from the field OFF shape of nine resting droplets, grouped by location and somite stage (ss). Tissue interfaces: olfactory placode/optic stalk (filled symbol) and retina/optic stalk (open symbol). Bar: median. (F) Kelvin–Voigt model used for the fits in G and H: the droplet’s capillary spring *k*_*d*_ in parallel with the tissue, a spring *k* in parallel with a dashpot *µ* (Fig. S3A,B). (G,H) Folded field ON phases of a droplet in the eye (G, blue) and of a droplet in the olfactory placode (H, green), averaged over ten cycles; shading, s.e.m. between cycles; dashed lines, single-exponential fits yielding timescales and viscosities indicated on the graphs.

We first analysed the resting shape of the droplets without magnetic field in order to measure the local anisotropic stresses *σ*_*T*_ using the interfacial tension measured against PBS (Fig. 2A–E). In the PSM, the droplets are deformed along the anteroposterior (AP) axis, consistent with previous results (Mongera et al., 2018), and *σ*_*T*_ ranges from 97 Pa to 488 Pa in three embryos of the same clutch at the 6–10 somite stage. In the head region, depending on the localisation of the droplet, we found different values and orientations for *σ*_*T*_, ranging from 33 Pa to 88 Pa in the eye (retina) and 86 Pa in the olfactory placode up to 201 and 134 Pa when droplets are located at tissue interfaces, between the olfactory placode and the optic stalk and between the retina and the optic stalk respectively (Fig. 2E). These values are in qualitative agreement with previous stress measurements carried out in zebrafish tissues using a similar approach (Mongera et al., 2018) or different types of sensors (Souchaud et al., 2022; Vasiljevic et al., 2026). Interestingly, the droplets at tissue interfaces are elongated along these interfaces, which could indicate the presence of compressive forces in these locations.

### Estimation of viscoelastic parameters

The viscoelastic parameters of a tissue can be measured by applying a step of magnetic field, which fixes the magnetic stress *σ*_*M*_, and observing the response of the tissue over time, which constitutes a so-called creep experiment. Importantly, the analysis of this experiment requires knowing the resting droplet deformation *ε*_off_ and its orientation. Indeed, the orientation of deformation is crucial in this calculation, which is why we use signed projections rather than absolute values. For instance, if the field is applied orthogonally to the resting elongation of a droplet, it begins by *rounding* the droplet, which means that the unsigned *ε* decreases. In such a situation, simply using the difference between the absolute values of *ε*_on_ and *ε*_off_ would return a negative modulus. This is why we always use the field-induced modulation Δ*ε*_∥_ of the signed projected strain in the following.

In the eye and in the olfactory placode, the field was applied in ten successive cycles, each made of a 120 s field ON phase followed by a 120 s field OFF phase. Because the field was applied in steps of the same duration and with the same field amplitude and direction, all deformation cycles could be aggregated into one curve for each tissue (Fig. 2G,H). The resulting deformation dynamics curves are well fitted by a single exponential saturation curve, indicating that the tissue behaves as a viscoelastic material leading to the Kelvin-Voigt model shown in Fig. 2F and further described in Fig. S3A,B. With this model, the creep response reads:

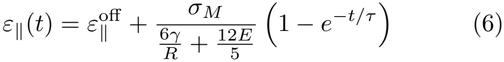

with 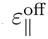 the resting deformation before the step, and

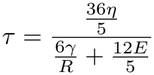

In concrete terms, the difference between the plateau of a creep curve and the resting deformation that precedes it (Fig. S3B) is

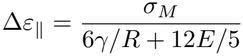

which returns the estimation of *E* given in Eq. (5), whatever the dynamics of the rise. Conversely, the fit of the whole curve using equation (6) yields the characteristic time *τ*, and the model then gives an estimate of the viscosity as 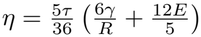 .

Averaging the estimated moduli over the ten cycles, we obtain 2.7 *±* 0.3 kPa in the eye and 3.3 *±* 0.3 kPa in the olfactory placode (mean *±* s.e.m. over the cycles of one droplet, Table 1). Fitting the folded rises yields *τ* = 44 s in the eye and 46 s in the olfactory placode (Fig. 2G,H), corresponding to viscosities of about 5 *×* 10^4^ Pa s and 6*×* 10^4^ Pa s respectively. At this stage both are neuroepithelial tissues of packed, adherent cells undergoing dynamic cell rearrangements (Sidhaye and Norden, 2017; Monnot et al., 2022; Breau et al., 2017; Aguillon et al., 2020; Bryan et al., 2020), which is compatible with the similar viscoelastic properties we report.

**TABLE 1.**
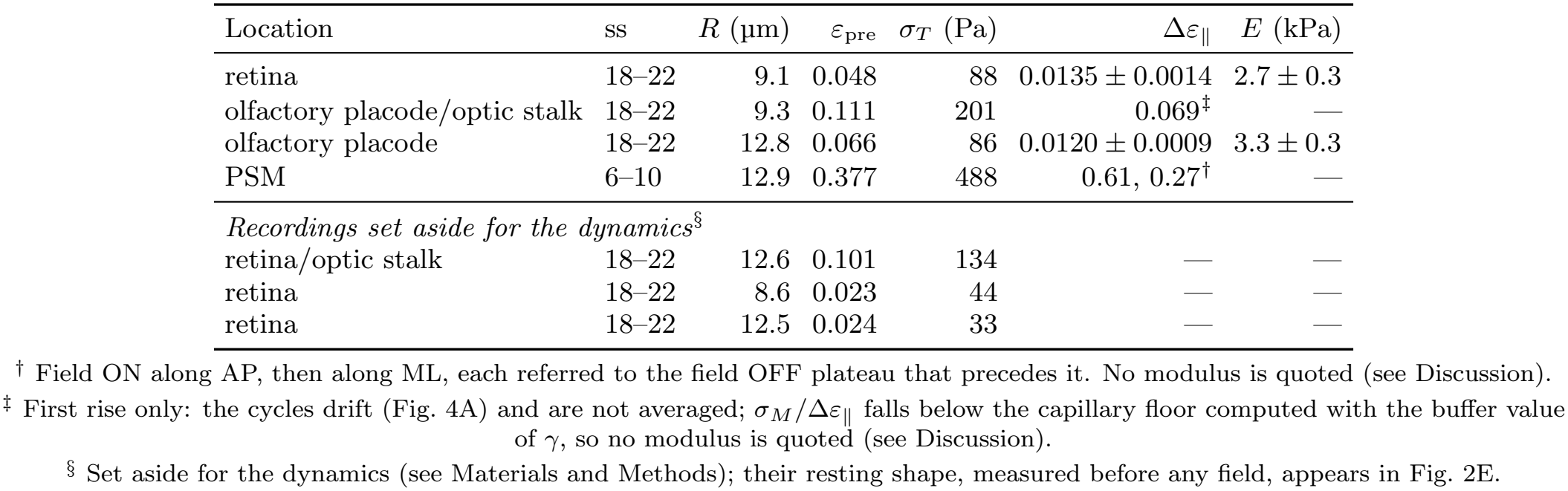
Droplet measurements in the zebrafish embryo. *R*, droplet rest radius; ss, somite stage; *ε*_pre_, deformation before any field; *σ*_*T*_ = 6*γε*_pre_*/R*, residual anisotropic stress computed with the buffer value of *γ* (Fig. 2E); Δ*ε*_∥_, field-induced modulation of the signed projected strain, each field ON plateau minus the field OFF plateau that precedes it, mean *±* s.e.m. over cycles; *E*, Young’s modulus from Eq. (5), mean *±* s.e.m. of the moduli of the individual cycles. All recordings at 1 frame s^*−*1^ and 160 mT.

| Location | ss | *R* (µm) | *ε*<sub>pre</sub> | *σ*<sub>*T*</sub> (Pa) | Δ*ε*<sub>∥</sub> | *E* (kPa) |
|---|---|---|---|---|---|---|
| retina | 18–22 | 9.1 | 0.048 | 88 | 0.0135 ± 0.0014 | 2.7 ± 0.3 |
| olfactory placode/optic stalk | 18–22 | 9.3 | 0.111 | 201 | 0.069<sup>‡</sup> | — |
| olfactory placode | 18–22 | 12.8 | 0.066 | 86 | 0.0120 ± 0.0009 | 3.3 ± 0.3 |
| PSM | 6–10 | 12.9 | 0.377 | 488 | 0.61, 0.27<sup>†</sup> | — |
| *Recordings set aside for the dynamics*<sup>§</sup> |  |  |  |  |  |  |
| retina/optic stalk | 18–22 | 12.6 | 0.101 | 134 | — | — |
| retina | 18–22 | 8.6 | 0.023 | 44 | — | — |
| retina | 18–22 | 12.5 | 0.024 | 33 | — | — |

<sup>†</sup> Field ON along AP, then along ML, each referred to the field OFF plateau that precedes it. No modulus is quoted (see Discussion).
<sup>‡</sup> First rise only: the cycles drift (Fig. 4A) and are not averaged; *σ*<sub>*M*</sub>/Δ*ε*<sub>∥</sub> falls below the capillary floor computed with the buffer value of *γ*, so no modulus is quoted (see Discussion).
<sup>§</sup> Set aside for the dynamics (see Materials and Methods); their resting shape, measured before any field, appears in Fig. 2E.

Note that no modulus is given for the PSM, where the deformation exceeds the limit set by the droplet’s own interfacial tension when the buffer value is used (see Discussion); instead this recording is used in the next section for its dynamics.

### Probing the relevant viscoelastic model

In developing tissues, the rheology can be more complex than the simple viscoelastic model identified in the previous section. In particular, a biological tissue can exhibit multiple characteristic times, corresponding to distinct biological processes. This type of analysis thus calls for fewer but longer field steps.

In the PSM, we performed experiments with a longer time window to probe a broader spectrum of relaxation times. Indeed, it has been shown that the tailbud region undergoes a liquid-to-solid transition along the AP axis (the tissue becoming more solid towards its anterior part) (Serwane et al., 2017; Mongera et al., 2023, 2018). Therefore, exploring the full range of timescales could be relevant in this tissue. Thus, two magnetic field steps were applied for 900 s along orthogonal axes and were each followed by field-free steps of 900 s (see Fig. 3A,B). In addition to this, the droplet’s anisotropic stress before any field application was also measured (see Fig. 3A) during 17 s.

**FIG. 3.**
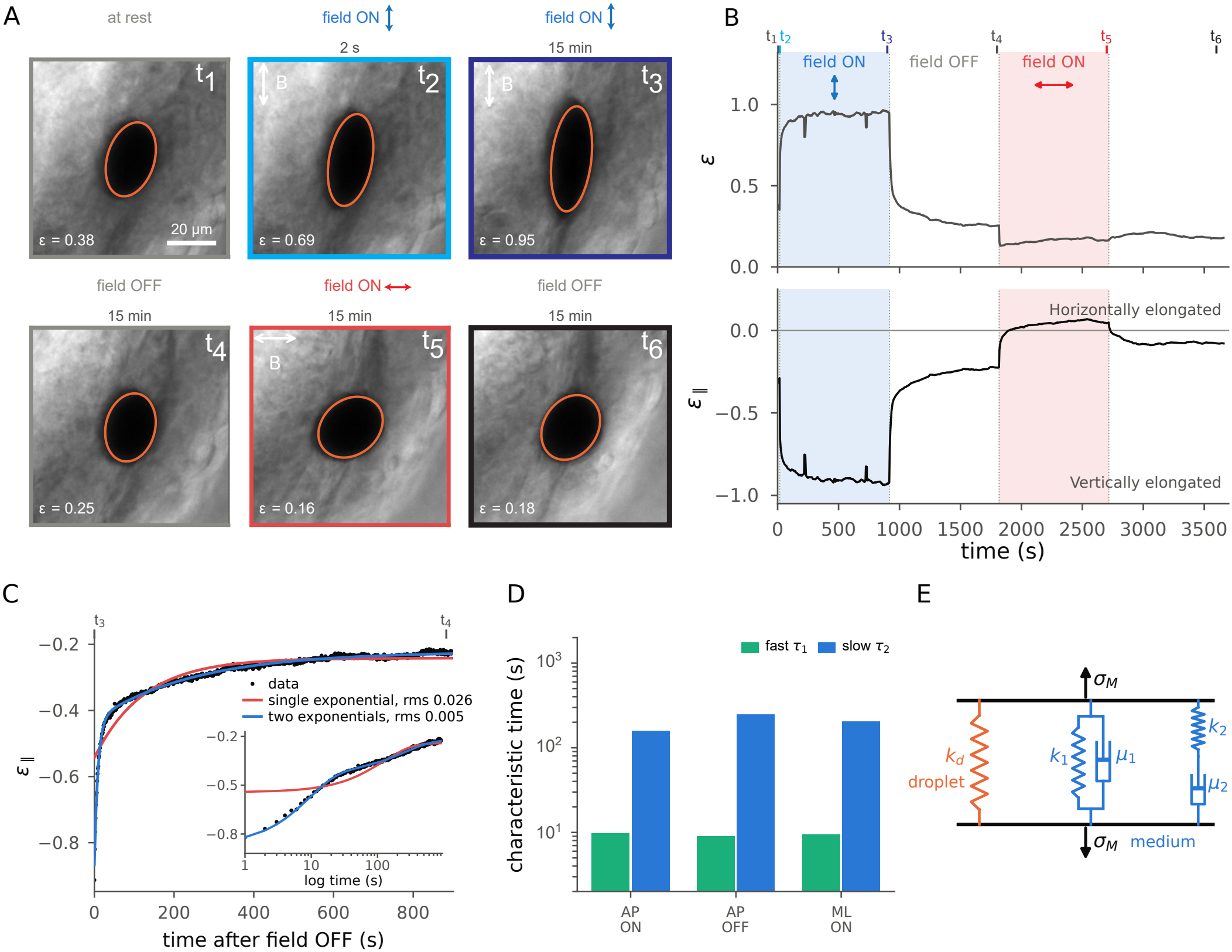
Two relaxation timescales describe mechanics in the PSM. (A) Bright-field images of a ferrofluid droplet at six points *t*_1_ to *t*_6_ of the protocol, reported on the time axis in panel (B). Orange line: sub-pixel ellipse fits. Frame colour: field direction as in (B). White arrows: applied field axis. (B) Droplet elongation over the field application protocol (total duration 61 min, acquisitions at 1 Hz): deformation *ε* (top) and its signed projection *ε*_∥_ (bottom). Shading marks field ON phases, coloured by field direction (blue, vertical; red, horizontal). Each phase lasts 900 s. (C) Field OFF relaxation of *ε*_∥_ (917 s to 1817 s) overlaid with single (red curve) and double exponential (blue curve) fits. The root-mean-square residuals (rms) calculated for each fit indicate a better agreement with a double exponential fit. Inset: same data on a logarithmic time axis, where the two timescales appear as two successive steps that a single exponential cannot follow. (D) Characteristic times obtained from the double exponential fits of three successive phases: field ON vertical, field OFF, field ON horizontal. The last relaxation phase is not analysed due to movements of the surrounding tissue. (E) Viscoelastic model with two timescales: the circuit of Fig. 2F with an additional Maxwell branch in parallel, a spring *k*_2_ in series with a dashpot *µ*_2_ (Fig. S3C,D).

We find that the first three phases of the protocol (field ON AP, field OFF, field ON ML) are better fitted with double exponential curves following the equation 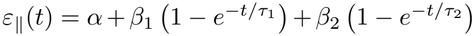, as shown in Fig. 3C. This allows for the estimation of characteristic times that yield successively *τ*_1_ = 9.7 s and *τ*_2_ = 160 s during the first field ON phase (AP axis), *τ*_1_ = 9.0 s and *τ*_2_ = 250 s during the first field OFF relaxation curve, 9.4 s and 206 s in the second field ON phase (ML axis) (Fig. 3D). Finding comparable timescales in three independent phases, with the field switched OFF or applied along two perpendicular directions, indicates that they reflect the tissue properties and not the fitting procedure. These two well-separated timescales are indicative of a viscoelastic model that is more complex than the one presented in Fig. S3A, used to derive Eq. (4). One way of introducing additional timescales in the rheological models is to add Maxwell branches (a spring in series with a dashpot) in parallel (Fig. 3E), each corresponding to an identified timescale (see further explanations in Fig. S3C,D). Interestingly, two distinct timescales have also been used to characterise the rheology of biological tissues in previous studies, both in spheroids (Yu et al., 2018) and in developing tissues (Serwane et al., 2017; Mongera et al., 2023).

### Probing tissue plasticity

Beyond timescales, the experiments performed in the PSM also shed light on local tissue plasticity. Indeed, after the field along the pre-stress direction, i.e. along the AP axis, was turned OFF, the droplet relaxed to its original deformation. However, when the field was applied orthogonally, in the ML axis, the droplet retained a residual deformation and did not return to its original shape. Irreversibility is thus oriented in this situation, suggesting that plasticity is more important in the ML axis in the PSM.

We tested plasticity with shorter cycles at the eye/placode interface. We applied the field in the axis orthogonal to the droplet’s resting elongation, so that it worked *across* the tissue–tissue boundary the droplet was lying along (Fig. 4). The field was applied along this axis in successive 2 min ON / 2 min OFF cycles. Strikingly, from the second field cycle onwards, the droplet no longer returned to its original elongation along the boundary but became more rounded instead. We interpret this as a signature of plasticity in the field direction, i.e. the droplet retained a residual deformation along the applied field axis which, together with the initial elongation orthogonal to it, amounts to an effective rounding. Interestingly, this effect accumulates over successive cycles, up until the point where the droplet remains constantly deformed along the field axis after 4 cycles. Over the same cycles, the response of the droplet to the field faded, from Δ*ε*_∥_ ≃ 0.07 and 0.12 on the first two cycles to 0.01 or less from the fifth. From the second cycle on, the magnetic stress brought the droplet within each field ON phase to about the same shape, nearly round or slightly elongated across the interface (*ε*_∥_ between 0 and 0.03). The droplet can thus be rounded up, but hardly elongated across the interface, as if the resistance of its surroundings rose sharply as soon as the droplet started to elongate across the interface. A stiffening of this region over the 40 min of the recording would also reduce the response. Again, these irreversible deformations indicate plasticity in a direction orthogonal to pre-stress in this area.

**FIG. 4.**
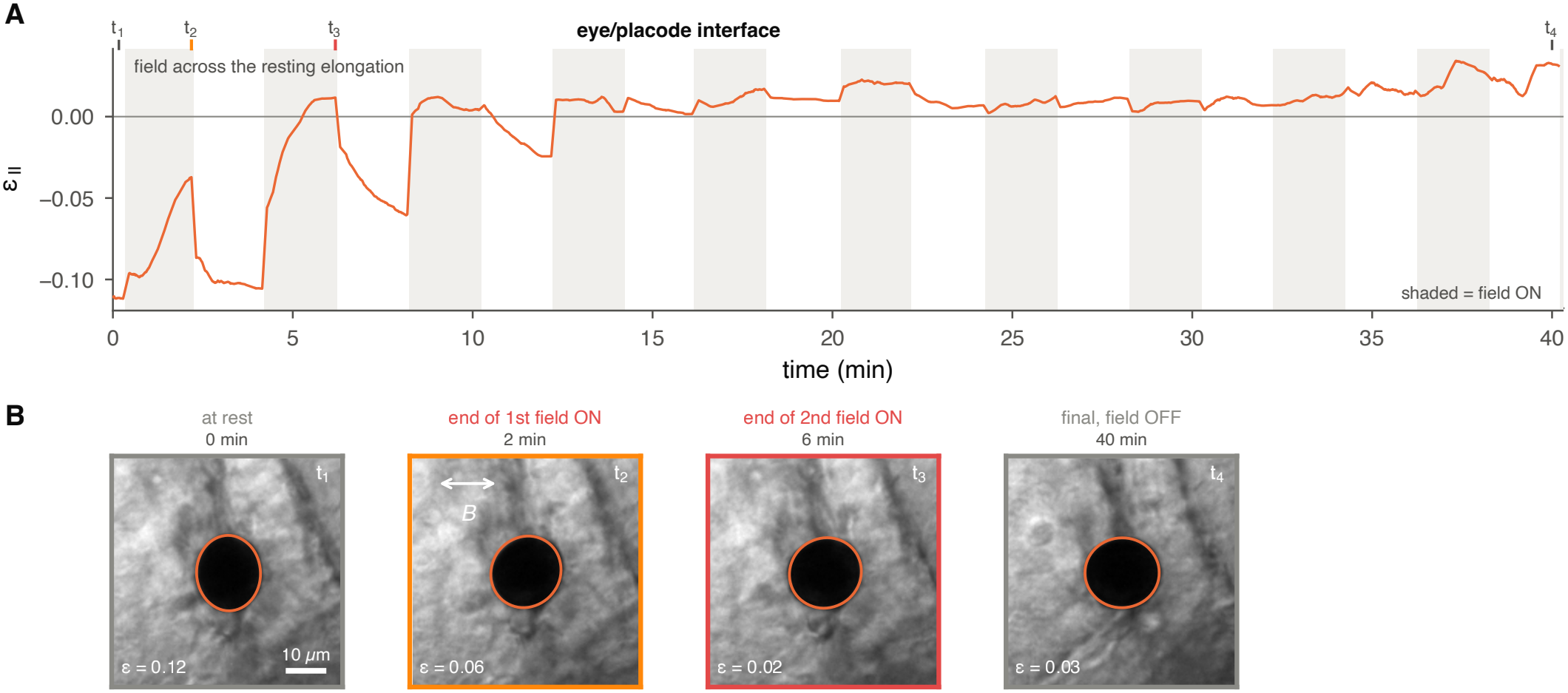
Probing plasticity at the eye/placode interface. (A) Signed projected strain *ε*_∥_ of a droplet located at the eye/placode interface over ten ON/OFF cycles with a magnetic field applied orthogonally to the interface, i.e. orthogonally to the droplet’s pre-stress (resting *ε*_∥_ ≃ *−*0.11). Shaded areas indicate field ON phases. Line, running mean over 20 s of the frames. Note that anatomical assignment of the field axis is not established for the head recordings. (B) Bright-field images of the ferrofluid droplet at four instants of the field application protocol, marked *t*_1_ to *t*_4_ on panel (A). Sub-pixel ellipse fits are overlaid on each image (orange). Field axis is indicated by a white arrow for field ON phases (framed in orange and red). Application of a magnetic field initially rounds up the droplet and eventually leads to an inversion of *ε*_∥_ because of accumulated irreversible deformations along the field axis.

Altogether these experiments indicate the importance of probing material properties along different axes in developing tissues. Indeed, plasticity is not necessarily isotropic, as revealed by experiments in the PSM.

## DISCUSSION

Morphogenesis depends on how tissues respond to the forces that shape them, and that response is set by stresses and material properties at a given time point. Therefore, these properties have to be measured locally and over the course of morphogenesis in order to understand the mechanics at stake. This can be done with the use of ferrofluid droplets that can report on local anisotropic stresses through their passive deformation, as well as on local material properties when they are used as micro-rheology probes under an applied magnetic field.

Generating a magnetic field with two nested, motorised Halbach rings turns the ferrofluid-droplet sensor into a rheometer that is portable and reconfigurable. The module mounts reversibly on a standard microscope stage, needs no power while imaging, and leaves every fluorescence channel free, since the droplet shape is measured on the bright-field contour with no fluorescent dye.

The most straightforward output is the *anisotropic stress*, read simply from the resting shape through capillarity, independent of the tissue modulus and of any assumption on the mechanical model for the tissue. The *plasticity* test comes next: it depends only on the shape the droplet returns to once the field is switched OFF.

The *characteristic times* are well determined once the field phases are long enough. The absolute *modulus* is the least well constrained, for a reason that Eq. (4) makes explicit: the droplet resists deformation through its own interfacial tension as much as through the tissue, and that self-resistance, ^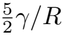^, is of the same order as the stiffness of a soft tissue; extracting *E* therefore means subtracting two numbers of similar magnitude, which propagates errors. A further limitation comes from imaging: the shape analysis uses the two-dimensional outline of the droplet. If a droplet is pre-stressed with an orientation that is tilted with respect to the optical axis, this 2D outline underestimates the elongation, and its principal axes are in general not the projections of the three-dimensional ones. Moreover, the field is applied in the imaging plane, thus imposing a deformation that is not aligned with the one set by the pre-stress. To overcome this issue, droplets that appeared tilted when scanning the focal plane in bright field were not recorded.

Every absolute stress given by this method is proportional to the interfacial tension between the droplet and the tissue *γ*, and every modulus depends on it through the capillary term of Eq. (5). The interfacial tension used for these calculations was estimated from pendant drop experiments in PBS buffer. Therefore the interpretation of anisotropic stresses can only be relative since the interfacial tension *in vivo* might differ from the one measured in PBS. Yet, this approach can be sufficient to decipher some biological processes, as done by Mongera et al. (2018). Interestingly, in the zebrafish embryo head tissues, the tissues appear stiffer than the posterior tissues in which ferrofluid droplets have been used before (Serwane et al., 2017; Mongera et al., 2023), and several explanations are plausible: these are different regions of the embryo, at a later stage of development and morphogenesis, when cells are more differentiated and cell densities and both cell–cell and cell–matrix adhesion are expected to be higher. The droplet deformations reported in the PSM clearly evidence that the interfacial tension in this tissue is lower than the one measured in the buffer. Indeed, the droplet’s own interfacial tension resists its deformation, so that under a given magnetic stress the droplet cannot deform beyond a maximum set by *γ*, even in a medium that offers no resistance at all; in Eq. (4), this means that *σ*_*M*_ */*Δ*ε*_∥_ can never fall below 6*γ/R*. In the PSM the droplet deforms well beyond that maximum, indicating that its interfacial tension must be lower than in a PBS buffer. This lowered tension can be due to an enrichment at the oil-water interface of a range of amphiphilic molecules that are present inside a biological tissue. As a consequence, the residual stresses in Fig. 2E are upper bounds and could be substantially smaller, while the moduli reported in Table 1 are lower bounds. However, if absolute values are necessary, the protocol used for the calibration in gels, which reports droplet deformation as a function of droplet size, can be performed within the tissue of interest in order to extract both the local interfacial tension and the Young’s modulus (Fig. 1G).

Two observations stand out in our measurements carried out in zebrafish embryos: in the PSM the droplet relaxes with two well-separated times; at the eye/placode interface the loading irreversibly modifies its resting shape. The two-timescale response in the PSM calls for a modelling of local tissue mechanics with two viscoelastic branches, associated with two timescales *τ*_1_ and *τ*_2_. Our observation of two well-separated timescales in the PSM is consistent with the two relaxation timescales reported in the zebrafish tailbud (Serwane et al., 2017; Mongera et al., 2023). Note that the dissipation behind these characteristic times can only come from the tissue, since the droplet’s own oil relaxes much faster; their values nonetheless also depend on the droplet, through its capillary stiffness 6*γ/R* (Fig. S3A).

In this framework, the faster timescale, on the order of 10 s, could be associated with the deformation of the cells themselves, which is set by the dynamics of the cytoskeleton, primarily the actomyosin cortex, with a possible contribution of microtubules and intermediate filaments. Conversely, the slower timescale, on the order of 200 s, likely represents the time associated with junctional remodelling for neighbour exchange under the load applied by the deforming droplet (Tlili et al., 2020; David et al., 2014; Mongera et al., 2023). These hypotheses can be tested with perturbative approaches: affecting intercellular adhesion strength or dynamics should displace the slow branch, while perturbing the cytoskeleton, for instance by inhibiting myosin II activity, should tune the short timescale. Droplet micro-rheology can thus address, *in vivo*, the cellular processes behind tissue mechanics, independently of any modulus determination.

The junctional remodelling dynamics are inherently linked to plasticity. For instance, if the cells exchange neighbours irreversibly under the mechanical perturbation imposed by droplet deformation, then the droplet cannot return to its initial shape. Such intercalation events are at the origin of the fluidisation in the zebrafish tailbud during axis elongation (Mongera et al., 2018, 2023). In addition, the properties of the extracellular matrix can also lead to plasticity signatures. For instance, at the eye/placode interface, where plasticity is the clearest, the droplet lies between the basement membranes that surround the two tissues (Monnot et al., 2022; Tignard et al., 2024; Bryan et al., 2020; Soans et al., 2022), and an irreversible remodelling of these matrix layers under load would leave exactly the signature we observe. Finally, plasticity could be envisioned at the scale of the cell itself (Molnar and Labouesse, 2021). Upon repeated contraction/relaxation cycles, the cell can keep irreversible deformations at each step akin to ratchet mechanisms (Muñoz and Albo, 2013; Lardennois et al., 2019). This could for instance explain the short timescale accumulation of plasticity observed in Fig. 4. Unravelling these distinct contributions to plasticity in developmental processes could typically be addressed by the combination of droplet micro-rheology with perturbative approaches.

In addition to all the measurements discussed above, our magnetic set-up design also offers the possibility to tune the amplitude of the field, which we did not explore experimentally. By rotating the rings gradually in opposite directions, the amplitude of the magnetic field can indeed be controlled continuously rather than in a step-wise manner (Fig. S1). This would allow us to identify the threshold stress beyond which the surrounding tissue deforms irreversibly, i.e. its yield stress. At the local scale, tuning the field amplitude also makes it possible to tune the load applied to cellular junctions. How T1 transitions are triggered as a function of this load would give the possibility to directly probe the energy barrier to cellular rearrangements (Popović et al., 2021; Sun et al., 2023) and to bridge the gap between junction mechanics and tissue-scale elasto-plasticity in a unified framework (Nguyen et al., 2025).

In summary, this easy to implement design, used with commercially available ferrofluids, gives access to a range of mechanical properties inside developing tissues with great modularity. As shown through the variety of experiments carried out in this study, simply tuning the field application protocol, for instance by using different time windows or field orientations, should help decipher a range of biological processes *in vivo*.

## MATERIALS AND METHODS

### Field generator

To create the Halbach rings, two sets of twelve magnets (3/16 and 3/8 inch cubes, K&J Magnetics B333-N52 and B666-N52) are pressed in custom 3D-printed housings, which are held concentric with two plastic slewing bearings with glass balls: one bearing attaches the small ring to the large one, the other attaches the large ring to the aluminium baseplate. Both rings are topped by a toothed profile and are driven by stepper motors through timing belts. The motors are driven by an Arduino Uno microcontroller running the grbl software and two stepper motor drivers. The baseplate is mounted to the microscope stage through vibration isolators. The axial field profile was measured with a Hall probe on the optical axis; in-plane values were measured at 1 cm from the centre, along and perpendicular to the nominal field direction, with the probe oriented along the field and then orthogonal to it. One centimetre from the centre, the field is 174 mT along the field direction and 167 mT perpendicular to it, symmetrically on both sides; extrapolating this quadratic variation, the field is uniform to 6 *×* 10^*−*3^ % over the field of view (494 *×* 395 µm) and to 2 *×* 10^*−*5^ % over a droplet. With the probe orthogonal to the nominal field direction we measured less than 2 mT everywhere, so the field direction is defined to better than 0.8^*°*^. With the two rings opposed, the residual field at the centre was 1.4 *±* 0.2 mT along the field of the outer ring and 0.8 *±* 0.2 mT orthogonal to it, which gives a residual magnetic stress of 0.6 Pa, less than 1 % of *σ*_*M*_ . The parts list, construction files and motor control script are available in the repository (see Data availability).

### Ferrofluid and magnetisation

Droplets were a commercial ferrofluid (FF-60, magnetshop.net) supplemented with Tween 20 (P9416, Sigma-Aldrich) at 20 mg mL^*−*1^. Its density, weighed in the laboratory, is 1.153 g cm^*−*3^. Magnetisation curves *M* (*B*) were measured by vibrating-sample magnetometry on the formulation, up to 850 mT. The applied magnetic stress is the Maxwell pressure at the droplet poles, *σ*_*M*_ = *µ*_0_*M* ^2^*/*2, evaluated at the working field: *M* = 1.28 *×* 10^4^ A m^*−*1^ at 160 mT gives *σ*_*M*_ = 103 Pa.

### Interfacial tension

Pendant drops of the formulation were formed from a 23-gauge needle into PBS. Contours were extracted at sub-pixel resolution from the maximum of the transmittance gradient, and the Young–Laplace profile was fitted by orthogonal-distance minimisation with the apex position, the apex radius, the Bond number and the tilt free; the needle width, measured frame by frame, provided the length standard. Edge and needle biases were calibrated on synthetic drops rendered with the same optics, which recover the true tension to 1.4 *±* 0.5 % for Worthington numbers above 0.4, the conditioning criterion of Berry et al. (2015). The reported value is the mean over drops. The largest term of the uncertainty is the density difference between the oil and the buffer, which we weighed by hand, closely followed by the scatter between drops. The scatter between drops (25 %) is much larger than within a drop (5.5 %), so it is a property of the interface and not of the fit. One fitted drop of the formulation is shown in Fig. S2A. The Tween 20 dilution series of Fig. S2B was measured on the same oil, with the same needle and into the same buffer; those drops were left to equilibrate for 30 min and their profiles fitted with the Pendent Drop plug-in of Fiji (Daerr and Mogne, 2016), each point being the mean over the last 5 min of several drops.

### Polyacrylamide gels

Gels of 6 % acrylamide and 0.06 % bis-acrylamide were prepared following Cresens et al. (2026): 10 mL of solution (1500 µL of 40 % acrylamide (A4058, Sigma-Aldrich), 290 µL of 2 % bis-acrylamide (M1533, Sigma-Aldrich) and 7935 µL of water, degassed by sonication on ice, then 250 µL of 10 % ammonium persulfate (A/P470/46, Fisher Scientific) and 25 µL of TEMED (T7024, Sigma-Aldrich)) were cast between the lid and the bottom of a Petri dish, trimmed of their outer 5 mm and swollen for at least 16 h in PBS without calcium and magnesium, the buffer of the pendant-drop measurements. We departed from that protocol in two respects: the gels were swollen at 22 ^*°*^C, the imaging temperature, rather than 37 ^*°*^C, and the whole plate was swollen before cutting. Droplets were injected with the same system as in embryos, into the swollen gel (gel 1), into the polymerised gel before swelling (gel 2), or into the setting gel just after casting, the dish being closed afterwards (gel 3). The bulk modulus of this composition, *E* = 0.90 *±* 0.11 kPa, is the value reported by Cresens et al. (2026) from oscillatory rheometry (*E* = 3*G*^*′*^ from a time sweep at 1 % strain and 1 Hz, at least three replicates).

### Zebrafish husbandry

Wild-type zebrafish embryos were obtained by natural spawning. Our experiments were performed in accordance with the European Directive 2010/63/EU on the protection of animals used for scientific purposes, and the French application decree Décret 2013-118. The fish facility has been approved by the French Service for animal protection and health, with the approval number B-75-05-25. The DAP number of the project is APAFIS #47427-2023121817395554 v6. Prior to imaging, embryos were staged according to developmental landmarks and somite number, following the developmental table of Kimmel et al. (1995).

### mRNA injections

mRNAs were synthesised from linearised vectors using the mMESSAGE mMACHINE SP6 transcription kit (Ambion). Wild-type zebrafish embryos were injected at the one-cell stage with 120 pg of gap-GFP mRNA to label cell membranes. The labelling of cell membranes was used to assess the potential toxicity of the ferrofluid and in experiments where the head region is targeted, and to precisely localise the ferrofluid droplet in embryonic tissues (Fig. 2D).

### Ferrofluid droplet injection in embryos

Embryos were mounted in 0.5 % low-melting agarose in E3 medium supplemented with tricaine at the 12-somite (15 hpf) stage for ferrofluid injection in the head region and at bud stage (10 hpf) for injection in the PSM. Injections were performed using a glass capillary without a filament (Harvard Apparatus, 30-0042) loaded with the ferrofluid solution and connected to a pressure control system (FemtoJet 4i, Eppendorf). Embryos and droplets started to be imaged and processed from around 2 hours after the mounting and 1 hour after the injection.

### Imaging

Images were recorded with a 40× water-immersion objective (Leica HC APO L 40x/0.80 W U-V-I) and a Teledyne FLIR Chameleon3 CM3-U3-13Y3M-CS camera. Bright-field image sequences were acquired at one frame per second, 1280 *×* 1024 pixels, with a spatial calibration of 0.386 µm per pixel. The droplet appears as a dark, near-circular disk on the translucent tissue and requires no fluorescent label. Field protocols were either 120 s ON / 120 s OFF cycling after an initial field-free delay, or the single-pass protocol with 900 s phases described in the Results; direction changes are stated per figure. Every change of field angle shakes the sample slightly, displacing it by up to two hundred pixels, which is why the analysis re-detects the droplet on every frame rather than tracking from a fixed crop.

### Analysis of the gel calibration

Gel images were analysed with the same contour definition as the embryo recordings (see Image analysis), adapted to still images of large droplets: the half-maximum level is taken relative to a polynomial fit of the local background, contour points cut by the frame edge or lying against a neighbouring object are discarded, and the ellipse is fitted by orthogonal distances, then refitted after rejecting points beyond five median absolute deviations. Contours are corrected for the radial distortion of the objective, calibrated on two stage micrometers (0.387 µm per pixel at the centre, magnification 0.3 % higher 600 px away). The two largest droplets did not fit in one frame and were measured on two overlapping frames registered on their common contour. Δ*ε*_∥_ is the field ON minus field OFF strain projected on the field axis of each session, measured from the orientation of the field-induced deformation of all droplets imaged in that session (5.8^*°*^, *−*3.7^*°*^ and 4.5^*°*^ from the image vertical). Its uncertainty, shown as error bars in Fig. 1G, is the quadrature sum of the statistical uncertainty of the ellipse fit, the half-range of Δ*ε*_∥_ over contour levels from 0.4 to 0.6, the size of the distortion correction and, for the two stitched droplets, the change when the registration uses one edge only; the robust refit changed none of the values. It amounts to 0.8 % to 10 % of Δ*ε*_∥_, much less than the scatter between droplets (*χ*^2^ = 527 for 11 degrees of freedom), so the fit of Fig. 1G is unweighted: uncertainties on *E* and *γ* are its standard errors, and resampling droplets within each gel gives 0.70 kPa to 0.86 kPa and 2.2 mN m^*−*1^ to 3.6 mN m^*−*1^ as 95 % intervals.

### Image analysis

All quantities were extracted with an open-source Python package (`ferro_analysis`). Images are read at full bit depth and normalised by the range of the data type rather than per frame. The droplet is detected as a dark, compact, near-circular connected component of a background-subtracted contrast map, retained on area and circularity criteria; it is re-detected on each frame inside a region of interest around its predicted position, with a fallback to the full frame. The shape is then measured on a tight window: a support mask is taken as the dark component nearest the window centre, with a morphological opening that severs thin connections to tissue striations, and the deformation is obtained from a subpixel ellipse fit to the half-maximum contrast contour restricted to that mask, choosing among contours the one of largest enclosed area. Orientation is measured separately, from the second-order intensity moments of the masked droplet, because moments stay stable on a nearcircular droplet exactly where a contour fit flips by 90^*°*^. The tracking occasionally loses the contour on an isolated frame, which displaces the projected strain by several tenths for that frame alone. Such frames are rejected before the relaxation fits, as a departure from a running median larger than six median absolute deviations, the first few frames of each phase being protected because they carry the transition. They are rare: four frames out of 900 on the phase most affected. The number of frames rejected is written out with each fit. We validated the measurement on synthetic ellipses matched to the real images in contrast, noise and edge sharpness, and passed through the same masked pipeline as the data. The ellipse fit is unbiased on both quantities that enter the analysis, below 3 *×* 10^*−*5^ on *ε* and 0.14 % on the volume-conserving equivalent radius, whereas the moment-based *ε* carries a size-dependent bias, which is why moments are used for the angle and the fit for the strain. The noise floor of the shape measurement is *σ*_*ε*_ = 0.045 *b*^*−*3*/*2^ with *b* the fitted semi-major axis in pixels, that is 0.02 % to 2.6 % of the measured *ε* over our range of droplet sizes; this, and not a pixel-scale guess, is what we report as the measurement error. The residual of the ellipse fit is recorded as a quality metric: a droplet free in a homogeneous medium is an ellipse, so a faceted contour signals contact with a wall, a cavity or a neighbouring droplet.

Field cycles are timed from the global frame-to-frame image motion, which spikes reproducibly at each field change; transitions are placed on a periodic comb and snapped to the rising edge of the shake, with a fallback to the nominal schedule. Frames shaken by a transition are excluded from the fits by an adaptive guard band. For each phase, the deformation is fitted to a single exponential, and all cycles of a given phase are also averaged on phase-relative time and fitted; the folded estimate is the one we report. The field-induced modulation Δ*ε*_∥_ is read cycle by cycle, as the plateau of each field ON phase, the median of its last 30 valid frames, minus that of the field OFF phase that precedes it, the rest before any field serving for the first cycle. Each cycle gives a modulus through Eq. (5), and the modulus we report is their mean, with its standard error. Two-exponential fits were performed phase by phase on the single-pass recording, which is not cycled and therefore not folded. Every run writes out the full time series, the fits, the figures and a record of the parameters used.

Of the recordings made with this module, four are analysed here (Table 1). The others were set aside either because the field protocol had a methodological bias (most often a field-free window too short to be read as a relaxation) or because the imaging session did not hold the droplet in focus and in frame for long enough to be exploited. The selection was made on the recording, not on the mechanical values it produced. Two further droplets in the PSM were imaged at rest only, on a single frame each; they are not listed in Table 1 and contribute the two lowest values of the PSM in Fig. 2E.

### Biocompatibility

All embryos followed through the experiments developed normally over the course of the recordings. Embryos ubiquitously expressing membrane-targeted GFP did not show any sign of cell death around the droplets (Fig. 2D).

## Use of AI tools

Claude was used to improve the image analysis scripts, starting from an existing code base written by the authors, and the preparation of the data deposit. The authors checked all code and its outputs, and take full responsibility for them.

## Data availability

The raw images, the magnetometry and field measurements, and the results derived from them are available at Zenodo (https://doi.org/10.5281/zenodo.24400524; Harmand et al., 2026b). The analysis code, which recomputes every number and figure panel of this article from these data, is available at Zenodo under the MIT licence (https://doi.org/10.5281/zenodo.24400756; Harmand et al., 2026a). The design files of the field module (3D models, files for printing, drawings and parts list) and its motor control script are available at Zenodo under the CERN Open Hardware Licence v2 – Permissive (https://doi.org/10.5281/zenodo.24400711; Harmand et al., 2026c).

## ACKNOWLEDGMENTS

The authors thank Delphine Talbot for her help with vibrating-sample magnetometry experiments. The authors also thank France Lam and Romain Naillon for their support at the IBPS imaging facility, and all the IBPS aquatic platform members for fish care. The authors thank Marion Baraban, Jonathan Fouchard, Alexis Prevost and Elie Wandersman for critical reading of the manuscript.

## FUNDING

This work was supported by the Agence Nationale de la Recherche (ANR-23-CE13-0025-02-MECAMATRIX to M.A.B. and L.-L.P.), the i-Bio international PhD Program (O.V.), the Fondation pour la Recherche Médicale (postdoctoral fellowship ARF202409019400 to C.C.), the Réseau EDEN-André Picard (seed grant to C.C.) and the Institut de Biologie Paris-Seine (seed grant to N.H. and C.C.).

## COMPETING INTERESTS

No competing interests declared.

## SUPPLEMENTARY FIGURES

**FIG. S1.**
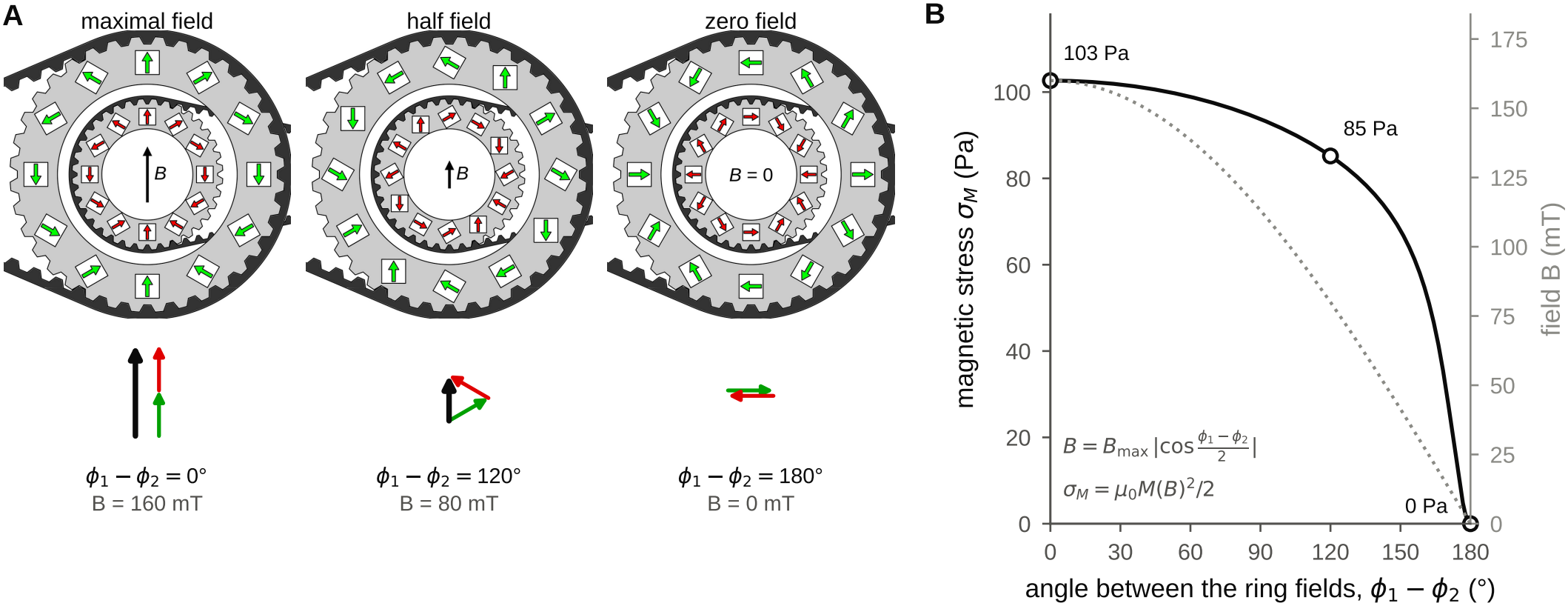
Tuning the field amplitude by counter-rotating the rings. (A) Two Halbach rings in three configurations: maximal field (ring fields aligned, *ϕ*_1_ *− ϕ*_2_ = 0), half field (*ϕ*_1_ *− ϕ*_2_ = 120^*°*^) and zero field (*ϕ*_1_ *− ϕ*_2_ = 180^*°*^). Green and red arrows on the magnets: magnetisation direction; black arrow in the bore, resulting field *B*. Bottom sketches: fields of the outer (green) and inner (red) rings added tip to tail, and their sum (black). (B) Magnetic stress *σ*_*M*_ at the centre as a function of the angle between two rings of equal strength, *B*_max_ = 160 mT, computed with *B* = *B*_max_ |cos((*ϕ*_1_ *− ϕ*_2_)*/*2) | (Eq. (1)) and the magnetisation curve of Fig. 1D. Dotted line: field *B* (right axis). Circles: the three configurations of (A).

**FIG. S2.**
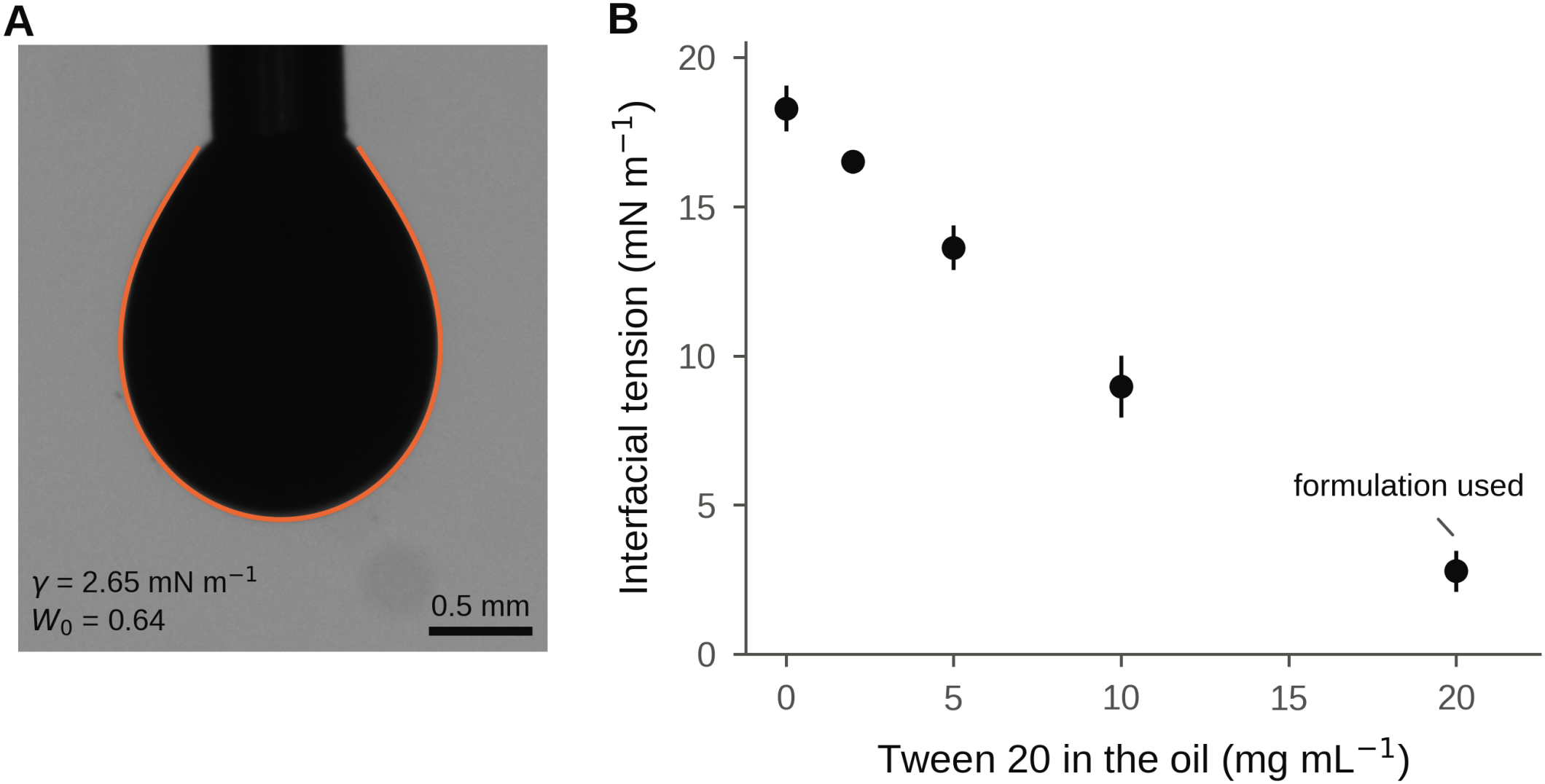
Interfacial tension measurement. (A) Image of a pendant drop of ferrofluid oil supplemented with 20 mg mL^*−*1^ Tween 20, in PBS, with the fitted Young–Laplace profile (orange). For this specific drop *γ* = 2.65 mN m^*−*1^ at a Worthington number of 0.64 (the mean over eight drops is 2.79 *±* 0.37 mN m^*−*1^). (B) Interfacial tension against PBS as a function of Tween 20 concentration in the oil. Error bars, dispersion between repeats at each concentration.

**FIG. S3.**
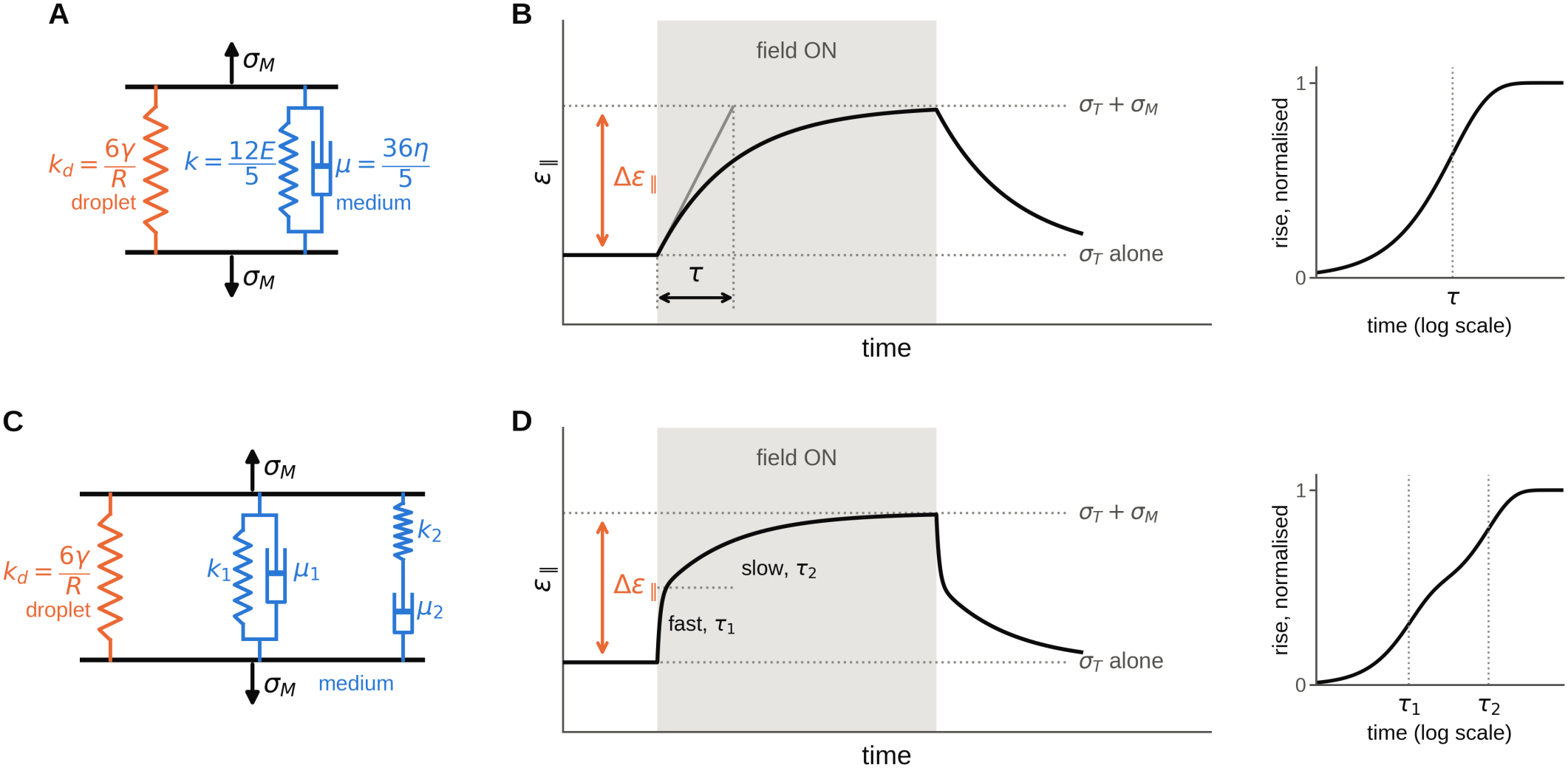
The two mechanical models used in this work. (A) One timescale: the droplet’s capillary spring, *k*_*d*_ = 6*γ/R* (orange), in parallel with the medium (blue), a spring *k* = 12*E/*5 in parallel with a dashpot *µ* = 36*η/*5 (Kelvin–Voigt element). (B) Response of this model to a field step. Before the field, the tissue stress *σ*_*T*_ alone sets the resting deformation; the field adds *σ*_*M*_, so that the difference Δ*ε*_∥_ gives *E* through Eq. (5). The rise has a single timescale, *τ* = *µ/*(*k*_*d*_ + *k*). (C) Two timescales: the same circuit with an additional Maxwell branch, a spring *k*_2_ in series with a dashpot *µ*_2_. (D) Response of this model: a fast rise (*τ*_1_), then a slow one (*τ*_2_), up to a plateau that does not depend on the dashpots, so that *E* is read as in (B). Right of B and D, the same rises on a logarithmic time axis: one step for one timescale, two steps for two (see the inset of Fig. 3C).

## Notes

### Competing Interest Statement

The authors have declared no competing interest.

